# The effector protein CgNLP1 of *Colletotrichum gloeosporioides* from *Hevea brasiliensis* disrupts nuclear localization of necrosis-induced transcription factor HbMYB8-like to suppress plant defense signaling

**DOI:** 10.1101/2022.01.23.477367

**Authors:** Guangyong Yang, Jie Yang, Qiwei Zhang, Wenfeng Wang, Liping Feng, Li Zhao, Bang An, Qiannan Wang, Chaozu He, Hongli Luo

**Affiliations:** Hainan Key Laboratory for Sustainable Utilization of Tropical Bioresources, College of Tropical Corps, Hainan University, Haikou 570228, China; Sanya Nanfan Research Institute of Hainan University, Hainan Yazhou Bay Seed Laboratory, Sanya, 572025, China

**Keywords:** *Colletotrichum gloeosporioides*, CgNLP1, pathogenic mechanism, *Hevea brasiliensis*, HbMYB8-like

## Abstract

- *Colletotrichum gloeosporioides* is the dominant causal agent of rubber tree anthracnose and leads to serious loss of natural rubber production. Fungi secrete numerous effectors to modulate host defense systems. Understanding the molecular mechanisms by which fungal effectors regulate plant defense is of great importance for the development of novel strategies for disease control.
- Here, we identified an NLP effector gene, *CgNLP1*, which contributed to virulence of *C. gloeosporioides* to rubber tree. Transient expression of CgNLP1 in the leaves of *Nicotiana benthamiana* induced ethylene production in plants. Ectopic expression of CgNLP1 in Arabidopsis significantly enhanced the resistance to *Botrytis cinerea* and *A. brassicicola*.
- CgNLP1 was shown to target a R2R3 type transcription factor HbMYB8-like in rubber tree, which localized on nucleus and induced necrosis in *N. benthamiana*. CgNLP1 disrupted nuclear accumulation of HbMYB8-like and suppressed necrosis induced by HbMYB8-like mediated SA signal pathway.
- This work suggested a strategy whereby *C. gloeosporioides* exploited CgNLP1 effector to suppress host defense to facilitate infection by disrupting the subcellular compartment of a host defense regulator HbMYB8-like.

## Introduction

*Colletotrichum* causes anthracnose on a wide variety of woody plants in tropical, subtropical and temperate climates (Liang et al., 2021). *C. gloeosporioides* is the dominant causal agent of rubber tree anthracnose and leads to serious loss of natural rubber production (Liu et al., 2018). To successfully infect and cause disease, phytopathogenic fungi need to form intimate associations and maintain constant communication with a susceptible host, and this communication can be achieved through the proteins, enzymes and metabolites secreted by phytopathogenic fungi (Heard et al., 2015). Effector proteins are small cysteine-rich proteins secreted by pathogens and play roles in virulence and the interaction between plant and pathogens. According to the innate immunity theory, plants have evolved two strategies to detect pathogens: one is the recognition of conserved microbial elicitors called pathogen associated molecular patterns (PAMPs) by receptor proteins called pattern recognition receptors (PRRs) on the external face of the host cell, which leads to PAMP-triggered immunity (PTI); another one is the recognition of pathogen virulence molecules called effectors by plant intracellular receptors called R protein, which leads to effector-triggered immunity (ETI) (Dodds & Rathjen, 2010).

The necrosis- and ethylene-inducing protein 1 (Nep1)-like proteins (NLPs) are an important effector family and named after the necrosis and ethylene-inducing protein (NEP1) firstly identified from culture filtrate of *Fusarium oxysporum* f.sp. *erythroxyli* (Bailey, 1995; Chen et al., 2018). NLPs widely distributed in oomycetes, fungi and bacteria, and shared a conserved necrosis-inducing phytophthora protein (NPP1) domain, typically containing a GHRHDWE heptapeptide motif which was crucial for toxicity (Gijzen & Nürnberger, 2006; Lenarčič et al., 2017). Based on the molecular structures and sequence comparison analysis, NLPs were divided into three phylogenetic group types: type I, type II and type III (Oome et al., 2014; Levin et al., 2019; Seidl & Ackerveken, 2019). Type I NLPs contained a conserved disulfide bond and were found predominately in plant microorganisms. Compared with type I, type II had a second conserved disulfide bond and an additional putative calcium-binding domain. Type III NLPs were different from the other types in the amino acid sequence of N- and C-terminal portion, and there is still very little experimental data available (Oome et al., 2014; Lenarčič et al., 2017). Based on the ability to induce necrosis, NLPs were divided into two groups (Schumacher et al., 2020). Group one was cytotoxic NLP proteins and group two was non-cytotoxic NLP proteins (Amsellem et al., 2002; Santhanam et al., 2013).

The members of group one as their names suggested were best known as a virulence factor for inducing necrosis and ethylene production in plant leaves (Bailey, 1995; Oome et al., 2014; Levin et al., 2019; Seidl &Van den Ackerveken, 2019). MoNEP1, MoNLP2 and MoNEP4, three NLP proteins from the hemibiotrophic plant pathogenic fungus *Magnaporthe grisea*, induced cell death and the production of reactive oxygen species in *N. benthamiana* (Fang et al., 2017). BcNEP1 and BcNEP2, two paralogous NLPs from the necrotrophic plant pathogenic fungus *Botrytis cinerea*, caused necrosis in all dicotyledonous plant species tested (Schouten et al., 2008). Normally, the cytotoxic NLP proteins were expressed during the switch from biotrophic to necrotrophic lifestyle in hemibiotrophic pathogens (Alkan et al., 2015). It had been proved that NLP-induced cell death was an active, light-dependent process that required HSP90 and interacts with a target site on extracytoplasmic side of dicot plant plasma membranes (Qutob et al., 2006). Biochemical analyses had revealed that the target of NLP on plant membrane was sphingolipid-glycosyl inositol phosphorylated ceramide (GIPC) which consisted of an inositol phosphoceramide and a head group consisting of glucuronic acid and a variable number and form of terminal hexoses. When NLPs bind to the head group of GIPC in monocotyledons, the three terminal hexoses prevented NLPs from inserting into the lipid bilayers of cell membranes, while the GIPC head of dicotyledons only had two terminal hexoses, allowing NLPs to insert into cell membranes and causing cell necrosis (Van den Ackerveken, 2017). Recent research results showed that a plant-derived LRR-only protein NTCD4 promotes NLP-triggered cell death and disease susceptibility by facilitating oligomerization of NLP in Arabidopsis (Chen et al., 2021).

Group two were non-cytotoxic NLP proteins which were often expressed during very early stages of the infection or keep at rather low levels during the whole course of infection (Cabral et al., 2012; Dong et al., 2012; Schumacher et al., 2020). These non-cytotoxic NLPs also acted as triggers of plant innate immune responses, including posttranslational activation of mitogen-activated protein kinase activity, deposition of callose, production of nitric oxide, reactive oxygen intermediates, ethylene, the phytoalexin camalexin and cell death (Fellbrich et al., 2000; Kanneganti et al., 2006; Qutob et al., 2006; Rauhut et al., 2009; Villela-Dias et al., 2014; Seidl & Van den Ackerveken, 2019). Ten different noncytotoxic NLPs (HaNLPs) from biotrophic downy mildew pathogen *Hyaloperonospora arabidopsidis* did not cause necrosis, but acted as potent activators of the plant immune system in *Arabidopsis thaliana*, and ectopic expression of HaNLP3 in Arabidopsis enhanced the resistance to *H. arabidopsidis* and activated the expression of a large set of defense-related genes (Oome et al., 2014). Moreover, it was also reported that multiple cytotoxic NLPs carried a motif of 20 amino acid residues (nlp20), and nlp20 could trigger PTI by binding in vivo to a tripartite complex RLP23-SOBIR1-BAK1 (Böhm et al., 2014; Albert et al., 2015). Ectopic expression of RLP23 in potato (*Solanum tuberosum*) enhanced immunity to *Phytophthora infestans* and *Sclerotinia sclerotiorum* (Albert et al., 2015).

In *Colletotrichum*, six NLP homologs were identified in the *C. higginsianum* genome. Of them, ChNLP1 induced cell death when expressed transiently in *Nicotiana benthamiana* and was expressed specifically at the switch from biotrophy to necrotrophy, whereas ChNLP3 lacks necrosis-inducing activity and was expressed in appressoria before penetration and (Kleemann et al., 2012). An effector NLP1 from *C. orbiculare* induced necrosis in *N. benthamiana* and also possessed MAMP sequence called nlp24 which triggered the ROS accumulation in leaf discs of Arabidopsis (Azmi et al., 2017). In this study, an NLP effector protein identified in *C. gloeosporioides* was named as CgNLP1 which contributed to virulence of *C. gloeosporioides* to rubber tree and enhanced the resistance of Arabidopsis to *B. cinerea*. A R2R3-type transcription factor HbMYB8-like, which localized on nucleus and induced necrosis, was identified as the target of CgNLP1 in rubber tree, and CgNLP1 disrupted the nuclear translocation of HbMYB8-like and inhibits HbMYB8-like-induced cell death mediated by SA aignling. Our results provide new insights into the molecular mechanisms of interaction between rubber tree and *C. gloeosporioides* mediated by CgNLP1.

## Materials and methods

### Biological materials and growth conditions

*Colletotrichum gloeosporioides* strain was isolated from the leaves of *Hevea brasiliensis* with anthracnose. *Botrytis cinerea* and *Alternaria brassicicola* courtesy of Tesfaye’s lab. All fungal strains were grown on potato dextrose agar (PDA) at 28°C in the dark. *Hevea brasiliensis* (Reyan 7-33-97) was grown on soil at 28°C. *Arabidopsis thaliana* columbia ecotype and *Nicotiana benthamiana* were grown on soil under fluorescent light (200 μE·m^2^·s^-1^) at 22°C with 60% RH and a 12-h-light/12-h-dark cycle.

### RNA Isolation, cDNA Synthesis, PCR amplification and qRT-PCR

Fungal total RNA was extracted using CTAB-LiCl method (Yang J et al., 2020). Plant total RNA were extracted according to the instruction of the polysaccharide polyphenol plant total RNA extraction kit (Tiangen: DP441). The contaminating DNA was eliminated using RNase-free DNase and the first-strand cDNA was synthesized using Revert Aid First Strand cDNA Synthesis Kit (Thermo Fisher). Quantitative RT-PCR analysis was performed using ChamQ Universal SYBR qPCR Master Mix (Vazyme Biotech: Q711-02) with the LightCycler 96 System (Roche). The *N. tabacum actin-7* (*NtActin 7*) and *H. brasiliensis* 18S rRNA (*Hb18S*) were used as an endogenous control for normalization. The primers used for quantitative RT-PCR and PCR amplification were list in Supplemental Table 1. Relative expression levels of target genes were estimated using the 2^-ΔΔCt^ method.

### Sequence analysis of CgNLP1 and HbMYB8-like

The amino acid sequences were deduced by DNAMAN software. Predictions of signal peptides were performed online by SignalP 5.0 analysis tool http://www.cbs.dtu.dk/services/SignalP/). The conserved domains were predicted using SMART website (http://smart.embl-heidelberg.de/). The multiple alignments of amino acid sequences were performed using ESPript 3.0 (http://espript.ibcp.fr/ESPript/cgi-bin/ESPript.cgi) and GeneDoc 2.7.0. The bootstrap neighbor-joining phylogenetic tree was constructed using Clustal X 2.0 and MEGA 7.0.

### Generation of *CgNLP1* knockout and complementary mutants

Based on the diagram of *CgNLP1* knockout vector, the 5’ and 3’ flanking region of *CgNLP1* were amplified from genomic DNA and ligated into the vector pCB1532 carrying the acetolactate synthase gene (SUR) cassette conferred resistance to chlorimuron ethyl (a sulfonylurea herbicide). For the complementation vector, the open read frame of *CgNLP1* fused with the 3 X FLAG coding sequence was cloned into the vector harboring the promoter of *ToxA*, the terminator of nos and the hygromycin phosphotransferase gene (HPH) (Figure S2a). Fungal transformation was carried out as described in our previous work (Wang et al., 2018). The mutants were analyzed by PCR analyses. The primers used for *CgNLP1*amplifing and mutant diagnosis were list in Supplemental Table 1.

### Ethylene production assays

The tobacco leaves transiently expressing *CgNLP1* was collected post Agrobacterium injection every 12 hours for 3 days and kept in sample bottles for 3 days. 0.5 mL of ethylene released from leaves then was moved to syringe for ethylene contents assayed using a portable ethylene analyzer (GC-FID 8890, Agilent, USA). Ethylene standards (99.99% purity) was used for standard curve construction.

### Construction of *CgNLP1* overexpression lines in Arabidopsis

The vector pER8-*CgNLP1*-FLAG was constructed with an estradiol-inducible promoter and then transformed into *Agrobacterium tumefaciens* GV3101. *Agrobacterium-mediated* flower dip method was used for Arabidopsis transformation of *CgNLP1*. T1 transgenic lines were screened with 30 mg/L hygromycin and Western Blot was used to confirm the positive transgenic lines after estradiol treatment. Appropriate total proteins from different lines were resolved on 10 % polyacrylamide gels and transferred to a nitrocellulose membrane (Bio-Rad). Anti-FLAG monoclonal antibody (1:1000, ab125243; Abcam) was used as primary antibody to detect the expression of CgNLP1 in transgenic lines. Horseradish peroxidase-conjugated anti-mouse antibody was used as the secondary antibody, and the signal was detected using the ECL western detection kit (RPN2232; GE Healthcare).

### Disease assays

For the pathogenicity test of *C. gloeosporioides* to rubber tree, Conidia were harvested from mycelium grown on PDA (potato dextrose agar, Difco) medium for 10 days in a 28°C incubator and resuspended in a solution of PD (potato dextrose broth, Difco) liquid medium to a final concentration of 2× 10^5^ conidia/mL. Then 10 μL of the conidial suspensions were inoculated onto the wounded rubber tree variety 7-33-97 leaves at “light green” stage. The inoculated leaves were kept in a moist chamber at 28 °C under natural illumination for 4 days, and the disease symptoms were scored.

*B. cinerea* and *A. brassicicola* disease assay were performed on detached leaves by drop inoculation. Both strains were cultured in 2×V8 solid medium and incubated at 22°C. Conidia of *B. cinerea* were collected and suspended in 1% Sabouraud maltose broth buffer (Difco) containing 0.05% (v/v) Tween-20 and the conidia density was adjusted to 2.5×10^5^ spores/mL before inoculation. Conidia of *A. brassicicola* were collected and suspended in water containing 0.05% (v/v) Tween-20 and the conidia density was adjusted to 5×10^5^ spores/mL before inoculation. In both cases, the droplets of 5 μL spore suspension above were inoculated on 4-week-old Arabidopsis leaves. The inoculated plants were kept under a transparent cover to maintain high humidity. The lesion diameters were measured to assess the levels of plant disease resistance. Each treatment contained three replicates of 9 leaves and the entire experiment was repeated three times.

### Subcellular localization and bimolecular fluorescence complementation (BiFC) assays

For Subcellular localization of HbMYB8-like, the coding sequence of *HbMYB8-like* was inserted into the transient expression vector 35S-MCS-mScarlet to generate recombinant plasmid HbMYB8-like-RFP. The vector MEIL-RFP was used as marker vector for plasma membrane and nuclear localization (Guy et al., 2013). For BiFC assay, the coding sequences of *CgNLP1* and *HbMYB8-like* were inserted into pSPYCE-YFP^C^ and pSPYNE-YFP^N^ respectively to generate recombinant plasmids pSPYCE-*CgNLP1*-YFP^C^ and pSPYNE-*HbMYB8-like*-YFP^N^. The above constructs were verified by sequencing and then introduced into Agrobacterium strain GV3101, respectively. The Agrobacterium carrying HbMYB-like-RFP was expressed in *Nicotiana benthamiana* leaf tissue by agroinfiltration for Subcellular localization. pSPYCE-*CgNLP1*-YFP^C^ and pSPYNE-*HbMYB8-like*-YFP^N^ was co-expressed in *Nicotiana benthamiana* leaf tissue for BiFC assay. The fluorescence distribution was observed with a laser confocal microscope (Leica TCS SP8).

### Phytohormones treatment

Seedings of Reyan7-33-97 were treated with 5 mM salicylic acid (SA), 1 mM methyl jasmonate (MeJA), 0.5 mM ethephon (ET) and placed in 25°C green house (Yang et al., 2020). After 0, 12, 24, and 48 hours following treating, leaves from seedings were collected and quickly frozen in liquid nitrogen and then stored in a −80°C refrigerator for RNA extraction. Two leaves from each seeding were picked and three seedlings were pooled as one biological replicate. Each seeding was harvested only once.

### Yeast Two-hybrid Screens

Yeast two-hybrid assays were performed with Matchmaker™ Gold Yeast Two-Hybrid System according to the manufacturer’s instructions (Clotech: No.630489). The coding sequence of *CgNLP1* was amplified and cloned into pGBKT7 to generate DNA binding domain bait protein fusion. The cDNA libraries of *Hevea brasiliensis* was constructed according to the manufacturer’s instructions (Clotech: PT4085-1). Interacting proteins were selected for selective medium lacking His, Leu, Trp and Ade. The putative interactors were then tested by assaying for the lacZ reporter gene activation as described in the Clotech protocol. The plasmids from the positive clones were then isolated and reintroduced into the original yeast bait and control bait strains to verify interaction.

### GST Pull-Down Assays

The Prokaryotic expression vectors pColdTM TF-CgNLP1 and pGEX-HbMYB8-like were constructed and transferred into *E. coli* BL21 (DE3) respectively. The expression of the fusion proteins was performed as described in the product manuals (Beyotime Biotechnology: P2262). The cell lysate supernatants containing GST-HbMYB8-like and His-CgNLP1 fusion protein were incubate with GST binding gels at 4°C for overnight, and the supernatants containing GST + His-CgNLP1 as the control. The pull-down reactions were analyzed by SDS-PAGE followed by Western Blot using anti-His (M20001; Abmart) and anti-GST (ab111947; Abcam) antibodies.

### Trypan blue staining

Four-week-old tobacco leaves were infiltrated with *Agrobacterium tumefaciens* GV3101 harboring empty vector pEGAD-eGFP and recombinant plasmid pEGAD-*CgCP1*-eGFP, respectively. Leaves were harvested 3h after infiltration and stained for cell death using Trypan Blue Staining Cell Viability Assay Kit (Beyotime Institute of Biotechnology, Haimen, China). For destaining, the leaf samples were boiled in bleaching solution (ethanol: acetic acid: glycerol=3:1:1) for 15 minutes.

### Statistical analysis

Statistical analysis was performed with IBM SPSS Statistics v.25. Differences at P < 0.05 were considered as significant.

## Results

### CgNLP1 was a type I NLP candidate effector

Based on the genome sequencing of *C. gloeosporioides* from *hevea brasiliensis*, the genes encoding extracellular secretory protein were predicted and one of them, named as *CgNLP1*, was amplified by RT-PCR. The open reading frame (ORF) of *CgNLP1* was 729 bp encoding a protein of 243 aa with two cysteine residues and a signal peptide (1-18aa) at its N-terminal (Figure S1). The amino acid sequences of CgNLP1 were aligned with some NLP proteins identified in fungus, oomycete and bacteria (Figure 1a). The alignment showed that CgNLP1 contained a typical NPP1 domain with a heptapeptide motif SHRHDWE, and had the highest homology with NLP protein from *verticillium dahliae*. The different types of NLP proteins identified in fungal species including CgNLP1 were used to generate a Neighbour joining tree (Figure 1b). Phylogenetic tree analysis revealed that CgNLP1 was clustered in the same branch with other type I NLP proteins. These results suggested that *CgNLP1* encoded a type I NLP effector protein.

**Fig. 1.**
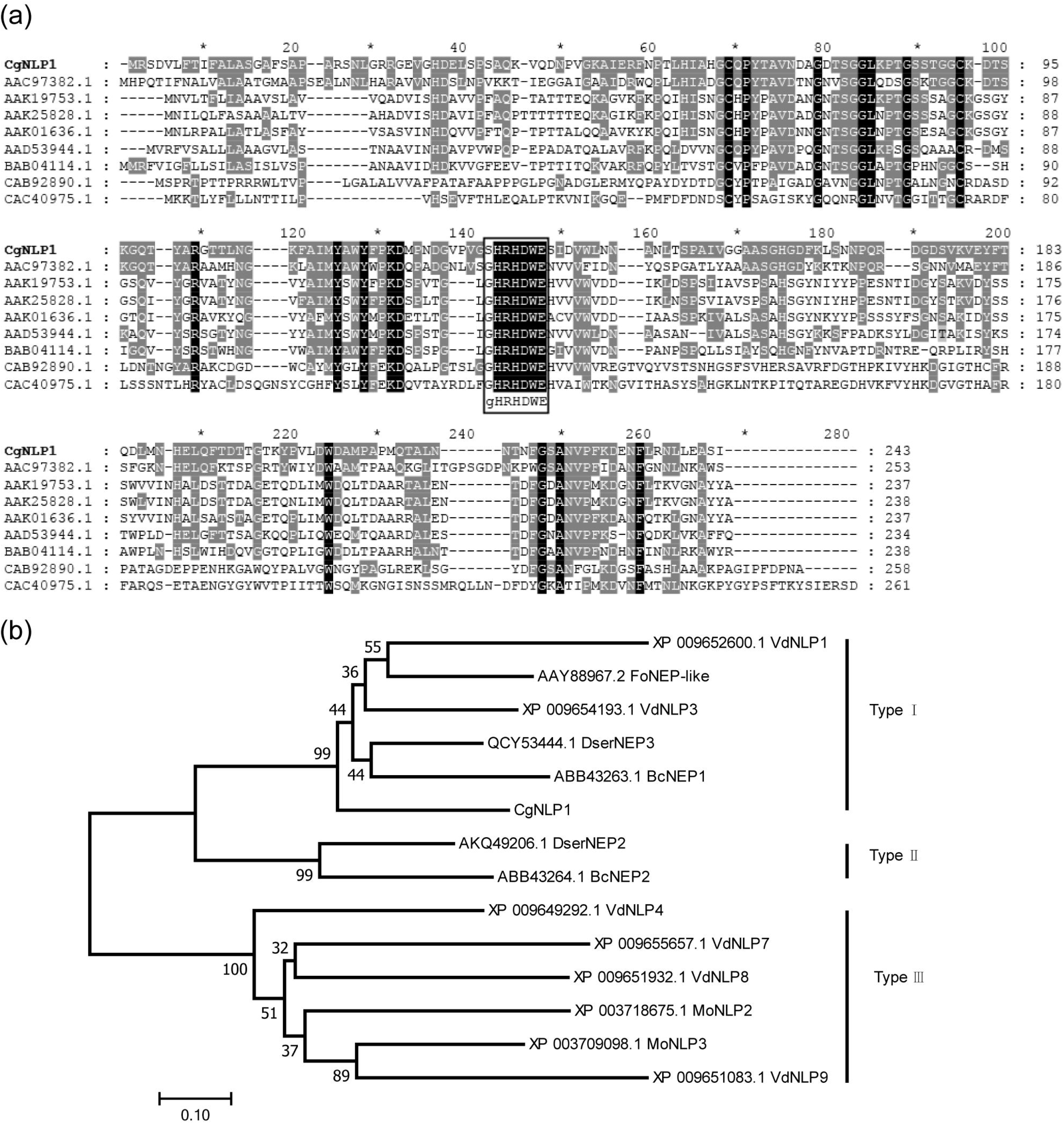
Multiple sequence alignment and Phylogenetic analysis of CgNLP1. (a) Alignment of the NLP proteins from different microorganism. Shading indicated regions of conservation in all (black), the same amino acid as CgNLP1 (grey) of sequences. NLPs used for alignment are from *Fusarium oxysporum* (AAC97382.1), *Pythium aphanidermatum* (AAD53944.1), *Phytophthora infestans* (AAK25828.1), *Phytophthora parasitica* (AAK19753.1), *Phytophthora sojae* (AAK01636.1), *Streptomyces coelicolor* A3(2) (CAB92890.1), *Alkalihalobacillus halodurans* C-125 (BAB04114.1) and *Vibrio pommerensis* (CAC40975.1). (b) Phylogenetic tree of CgNLP1 with different type of NLPs in fungi. VdNLP1 and VdNLP3 are from *Verticillium dahlia*, DserNEP3 is from *Diplodia seriata,* FoNEP-like is from *Fusarium oxysporum*, BcNEP1 and BcNEP2 are from *Botrytis cinerea*, DserNEP2 is from *Diplodia seriata*, PcNPP1 is from *Phytophthora cinnamomi*, MoNLP2 and MoNLP3 are from Pyricularia oryzae, VdNLP4, VdNLP7, VdNLP8 and VdNLP9 are from *Verticillium dahlia*.

### CgNLP1 contributed to the virulence of *C. gloeosporioides* to rubber tree

The *CgNLP1* knockout mutant (Δ*CgNLP1*) was obtained through gene homologous recombination technology and its complementary strain (Res-Δ*CgNLP1*) was generated by introducing *CgNLP1* into Δ*CgNLP1* (Figure S2). The detached leaf inoculation assay of Δ*CgNLP1* and Res-Δ*CgNLP1* on rubber tree was performed to explore the contribution of *CgNLP1* to the pathogenicity of *C. gloeosporioides*. Results showed that typical necrotic lesions were observed in the leaves inoculated with WT, Δ*CgNLP1* and Res-Δ*CgNLP1* (Figure 2a). At 4 dpi, the lesion size caused by Δ*CgNLP1* was significant smaller than that caused by WT, and there was no significant difference in the lesion size caused by Res-Δ*CgNLP1* compared with the WT (Figure 2b). These results indicated that the loss of *CgNLP1* resulted in reduced pathogenicity of *C. gloeosporioides*, suggesting the contribution of *CgNLP1* to the virulence of *C. gloeosporioides* to rubber tree.

**Fig. 2.**
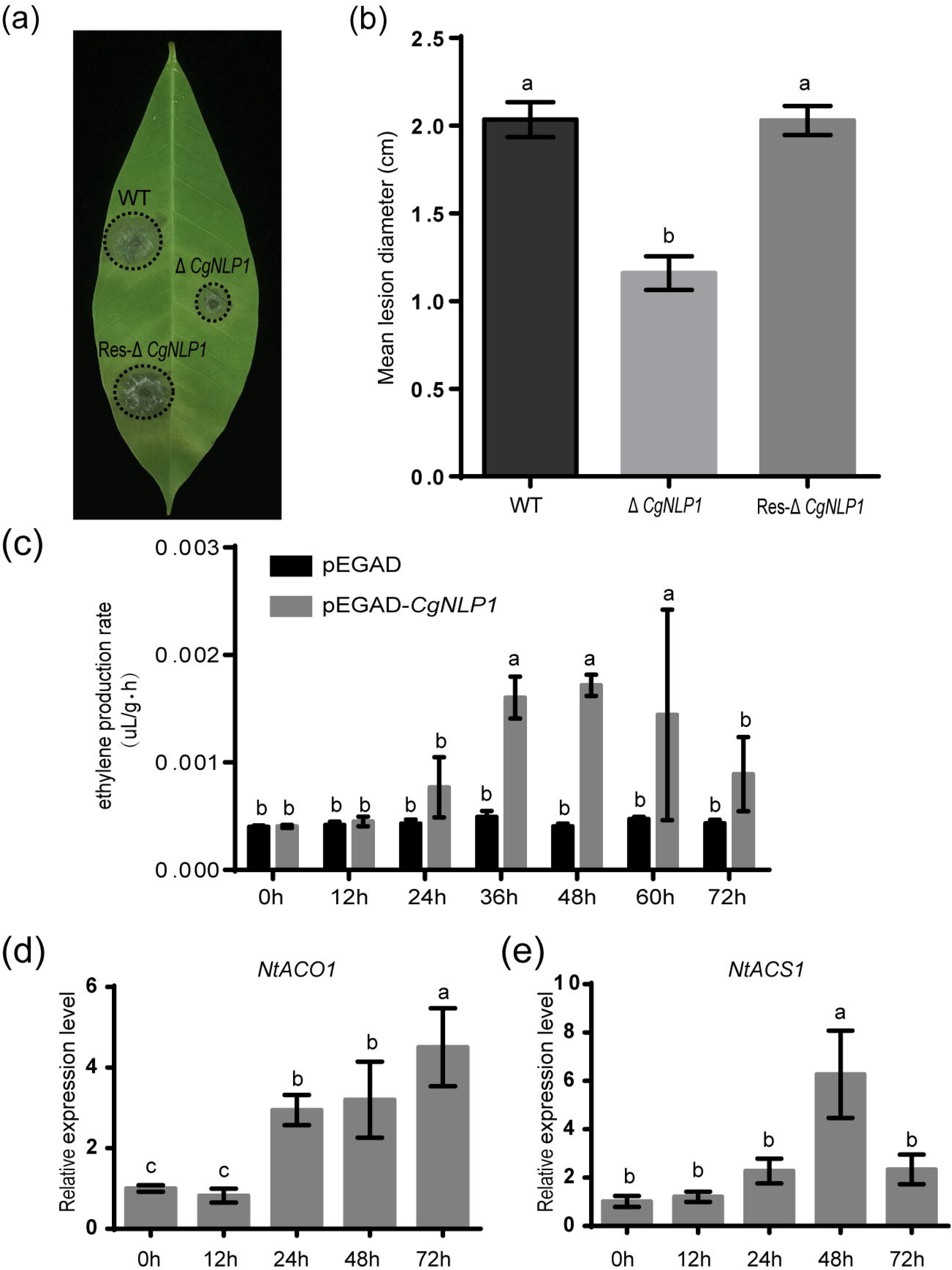
CgNLP1 contributed to the virulence of *C. gloeosporioides* to rubber tree and induced ethylene production in plants. (a) Disease Symptoms on rubber tree leaves at 4 days post inoculated with conidia of Δ*CgNLP1*, Res-Δ*CgNLP1* and WT. (b) Statistic analysis of lesion diameter after inoculation with WT, Δ*CgNLP1* and Res-ΔCgNLP1. (c) The ethylene content in tobacco leaves expressing *CgNLP1.* pEGAD represented the tobacco leaves expressing empty vector pEGAD and pEGAD-CgNLP1 represented the tobacco leaves expressing constructive vector pEGAD-CgNLP1. (d) The expression pattern of *NtACO1* (LOC107781126) in tobacco leaves expressing CgNLP1. (e) The expression pattern of *NtACS1* (LOC107831434) in tobacco leaves expressing CgNLP1 at different time points. Data are shown as the means ± SD from three independent experiments and columns with different letters indicate significant difference (P < 0.05).

### CgNLP1 induced ethylene production but not cell death

To determine the necrosis and ethylene inducing ability of CgNLP1 in plants, tissue necrosis observation and the ethylene content determination were performed in the tobacco leaves transiently expressing *CgNLP1* by agroinfiltration. No obvious tissue necrosis was observed in the infiltration area of tobacco leaves within 3 days post infiltration and trypan blue staining results showed no obvious difference between the leaves infiltrated with *A. tumefaciens* GV3101 harboring *CgCP1* gene and empty vector (Figure S3). The ethylene content in the leaves expressing CgCP1 was significantly higher than that in the leaves expressing empty vector (Figure 2c), and the expression levels of ACO and ACS, two key enzymes of ethylene synthesis, were significantly higher in leaves expressing NLP than that in control (Figure 2d and 2e). These data suggested that CgNLP1 induced ethylene production in plant but not necrosis and cell death.

### Ectopic expression of *CgNLP1* enhanced plant disease resistance

*CgNLP1* transgenic Arabidopsis plants driven by estradiol-induced promoter were generated to explore the roles of *CgNLP1* on plant disease resistance (Figure S4). We next studied the possible function of CgNLP1 in resistance against a fungus pathogen by analyzing disease phenotypes of the different CgNLP1 lines to *B. cinerea* and *A. brassicicola* inoculation. Four-week-old plants were challenged with a normally virulent strain of *B. cinerea* and *A. brassicicola* overexpression of CgNLP1 resulted in increased growth of the fungus and enhanced development of disease symptoms and that disruption of CgNLP1 led to decreased growth of the fungus and reduced development of disease symptoms (Figure 3a, 3b and 3c). Therefore, the Arabidopsis plants overexpressing *CgNLP1* significantly enhanced the resistance to *B. cinerea* and *A. brassicicola*, these results implied that exogenous expression of *CgNLP1* could improve plant disease resistance

**Fig. 3.**
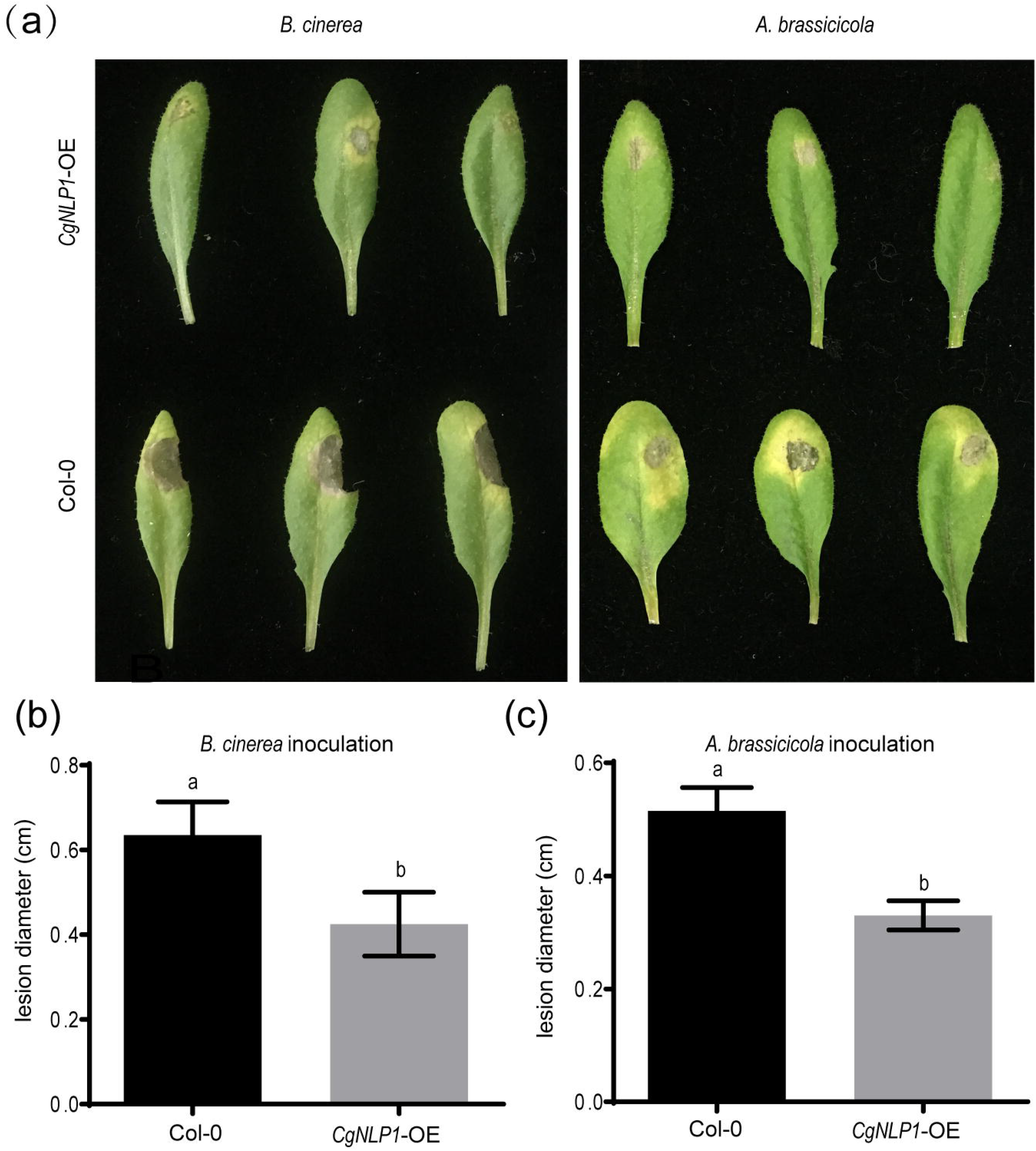
Overexpression of *CgNLP1* in Arabidopsis enhanced the resistance to *B. cinerea* and *A. brassicicola*. (a) Disease Symptoms on Arabidopsis lines overexpressing *CgNLP1* (*CgNLP1-OE*) and wild type (Col-0) at 3 days post inoculated with *B. cinerea* and *A. brassicicola*, respectively. (b) Statistic analysis of lesion diameter on CgNLP1-OE and *Col-0* after inoculation with *B. cinerea* and *A. brassicicola*. Data are shown as the means ± SD from three independent experiments and columns with different letters indicate significant difference (P < 0.05).

### CgNLP1 targeted rubber tree transcription factor HbMYB8-like

In order to elucidate the pathogenic mechanism of *C. gloeosporioides* on rubber tree, CgNLP1-associated proteins were identified from a cDNA library of rubber tree leaves by yeast two-hybrid screening using the full-length CgNLP1 as bait. After initial screening, 25 positive clones were sequenced and five of them were chosen for candidates. After verification, it is found that four of them had strong self-activation and only one of them, named as HbMYB8-like, which self-activation could be inhibited by adding an appropriate concentration of Aba. The *HbMYB8-like* gene was amplified by RT-PCR and verified by sequencing. The result showed that the full-length cDNA of *HbMYB8-like* gene was 1183 bp with a 903 bp ORF encoding 300 amino acids. HbMYB8-like protein contained two SANT domains which were typical features of MYB transcription factors. The results of multiple sequence alignment (Figure S5) and phylogenetic tree (Figure S6) showed that HbMYB8-like protein had typic adjacent repeats R2R3 and clusters with R2R3-MYB type MYB transcription factors from other plants, suggesting that HbMYB8-like belonged to the R2R3 type MYB transcription factors.

The interaction of CgNLP1 and full length HbMYB8-like was preliminarily verified through yeast two-hybrid (Figure 4a). Then Pull-down and BiFC assay were performed to further verify the interaction between CgNLP1 and HbMYB8-like in vitro. In pull down assay, His-tagged CgNLP1 and GST-tagged HbMYB8-like protein were expressed in *Escherichia coli* BL21 (DE3) respectively. The supernatant containing GST-tagged HbMYB8-like proteins were precipitated using anti-GST beads after co-incubation with supernatants containing His-tagged CgNLP1 and His protein only respectively, and the precipitates were detected using anti-His and anti-GST antibodies. It was found that only His-tagged CgNLP1could be pulled down by GST-tagged HbMYB8-like protein (Figure 4b). In BiFC assay, CgNLP1 was translationally fused with the C-terminal portion of YFP (CgNLP1-cYFP), and HbMYB8-like was fused with the N-terminal portion of YFP (HbMYB8-like -nYFP). CgNLP1-cYFP and HbMYB8-like -nYFP were introduced into *A. tumefaciens* and co-infiltrated into *N. benthamiana* leaves. Microscopic examination revealed YFP fluorescence only when the two constructs were co-expressed. Leaves from plants infiltrated with either of the constructs alone or in combination with the empty vector showed no fluorescence (Figure 4c). These results demonstrated that CgNLP1 interacted with HbMYB8-like.

**Fig. 4.**
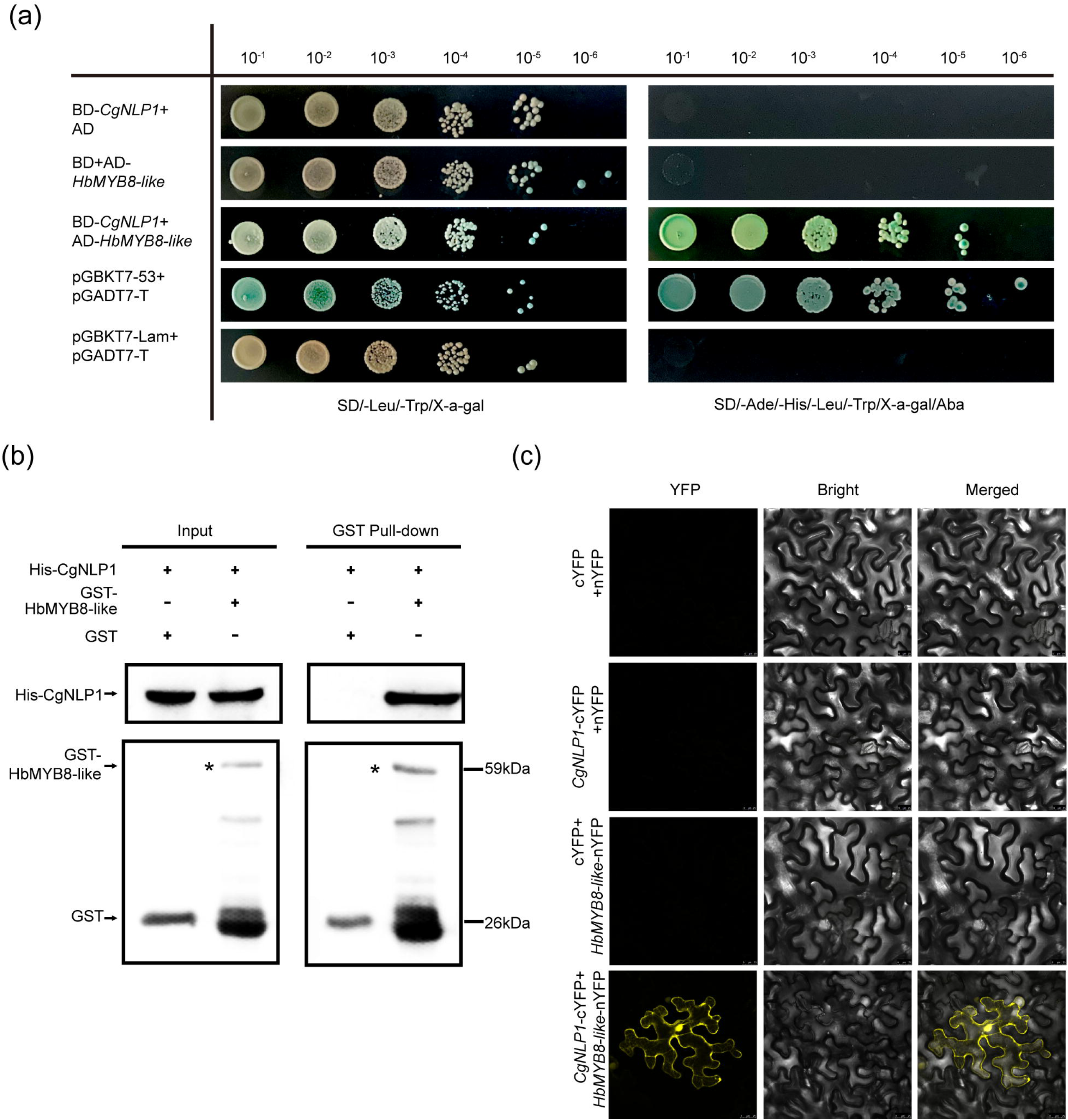
Screening and verification of the interaction between CgNLP1 and HbMYB8-like. (a) Verification of the interaction between CgNLP1 and HbMYB8-like by yeast two-hybrid assay. AD indicates pGADT7, BD indicates pGBKT7, pGADT7-T + pGBKT7-53 indicates a positive control, and pGBKT7-Lam+pGADT7-T indicates the negative control. SD/-Leu/-Trp/X-α-gal indicates the medium with X-α-gal, but lacking Leu and Trp. SD/-Ade/-His/-Leu/-Trp/Xa-gal/Aba indicates the medium with X-α-gal and Aba, but lacking Ade, His, Leu and Trp. 10^-1^, 10^-2^, 10^-3^, 10^-4^, 10^-5^, and 10^-6^ respectively refer to the yeast suspension with the initial OD value of 2.0 being diluted in a 10-fold gradient. (b) Verification of the interaction between CgNLP1 and HbMYB8-like by GST pull-down assay. His and GST stand for pCold™ TF and pGEX respectively. Arrows indicate GST- and His-tagged proteins. (c) Verification of the interaction between CgNLP1 and HbMYB8-like by bimolecular fluorescence complementation (BiFC) assay. cYFP+nYFP, *CgNLP1*-cYFP+nYFP, *cYFP+HbMYB8-like*-nYFP were used as a negative control. Scale bar□=□25um.

### HbMYB8-like protein localized on nucleus and induced necrosis

HbMYB8-like-RFP fusion protein was transiently expressed in *N. benthamiana* by agroinfiltration to determine the subcellular localization and explore the function on defense response. In the leaf tissues expressing only RFP, red fluorescence was observed throughout the entire cells. However, in the tissues expressing HbMYB8-like-RFP fusion proteins, red fluorescence was observed only on the nucleus (Figure 5a). After 2 days post infiltration with *A. tumefaciens* harboring *HbMYB8-like-RFP* gene, significant necrosis was observed in the infiltration area of tobacco leaf, while no necrosis was observed on the leaf infiltrated with *A. tumefaciens* harboring only RFP gene (Figure 5b).

**Fig. 5.**
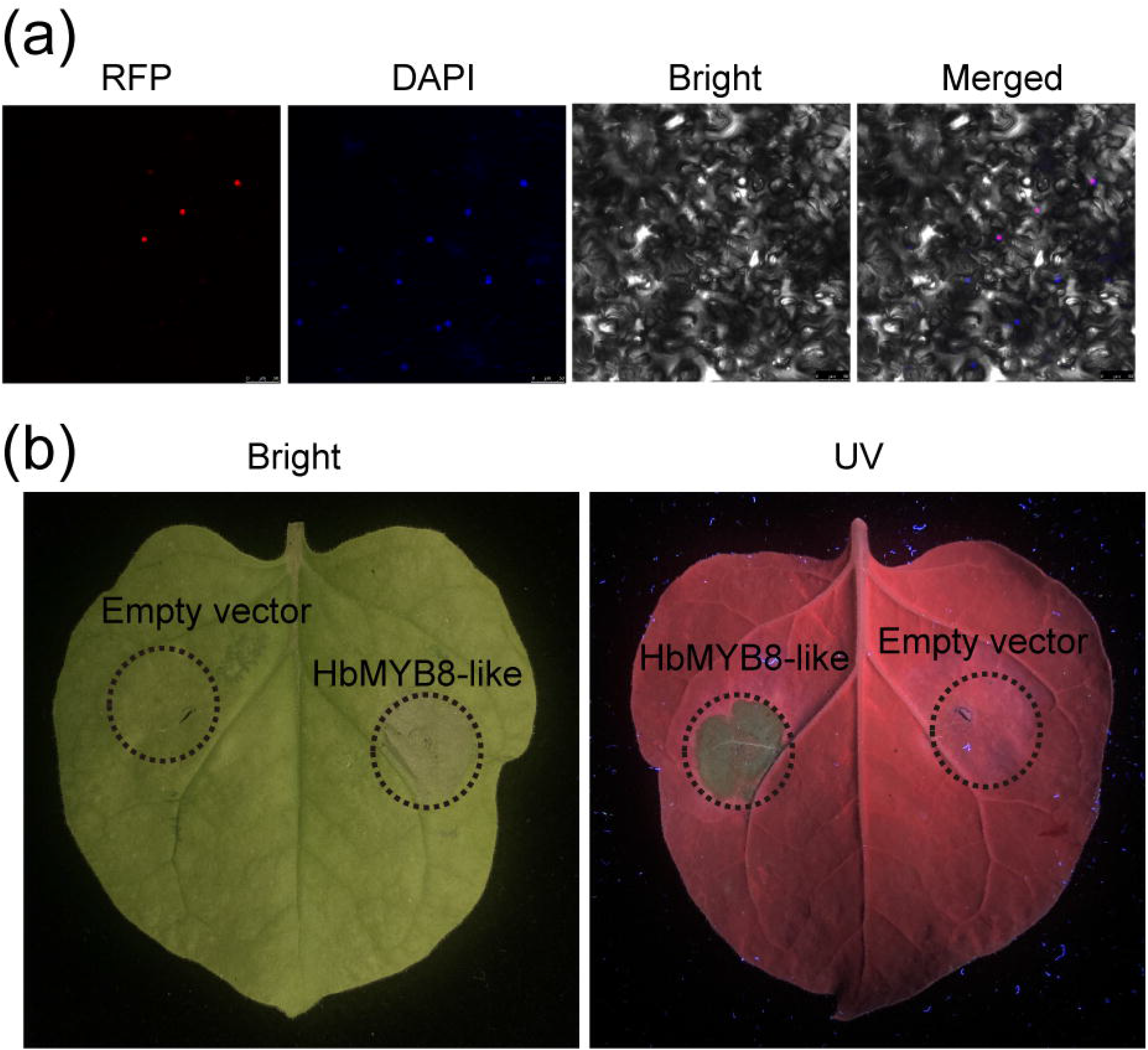
HbMYB8-like localized on nucleus and induced necrotic cell death. (a) Subcellular localization of HbMYB8-like protein in tobacco leaves. (b) Cellular necrosis induced by HbMYB8-like in tobacco leaves. Leaves were photographed under UV illumination (right) and normal light (left).

### *HbMYB8-like* responded to fungal phytopathogens and phytohormones in rubber tree

To explore the possible roles of *HbMYB8-like* in disease resistance of rubber tree, the expression profiles of *HbMYB8-like* were investigated in responding to *C. gloeosporioides* and *Erysiphe quercicola* which caused rubber tree anthracnose and powdery mildew respectively. In the rubber tree leaves inoculated with *C. gloeosporioides* and *E. quercicola*, the expression of *HbMYB8-like* was upregulated significantly at 24 hr post inoculation (hpi) (Figure 6a-b). In addition, the expression profiles of *HbMYB8-like* responding to different phytohormones were also investigated. The results showed that the expression of *HbMYB8-like* was significantly induced more than 8 times at 12 hours after SA treatment, and was significantly down-regulated by JA and ET treatments (Figure 6c-e). These results suggested that *HbMYB8-like* was involved in the resistance to fungal phytopathogens through SA, JA and ET mediated signaling in rubber tree.

**Fig. 6.**
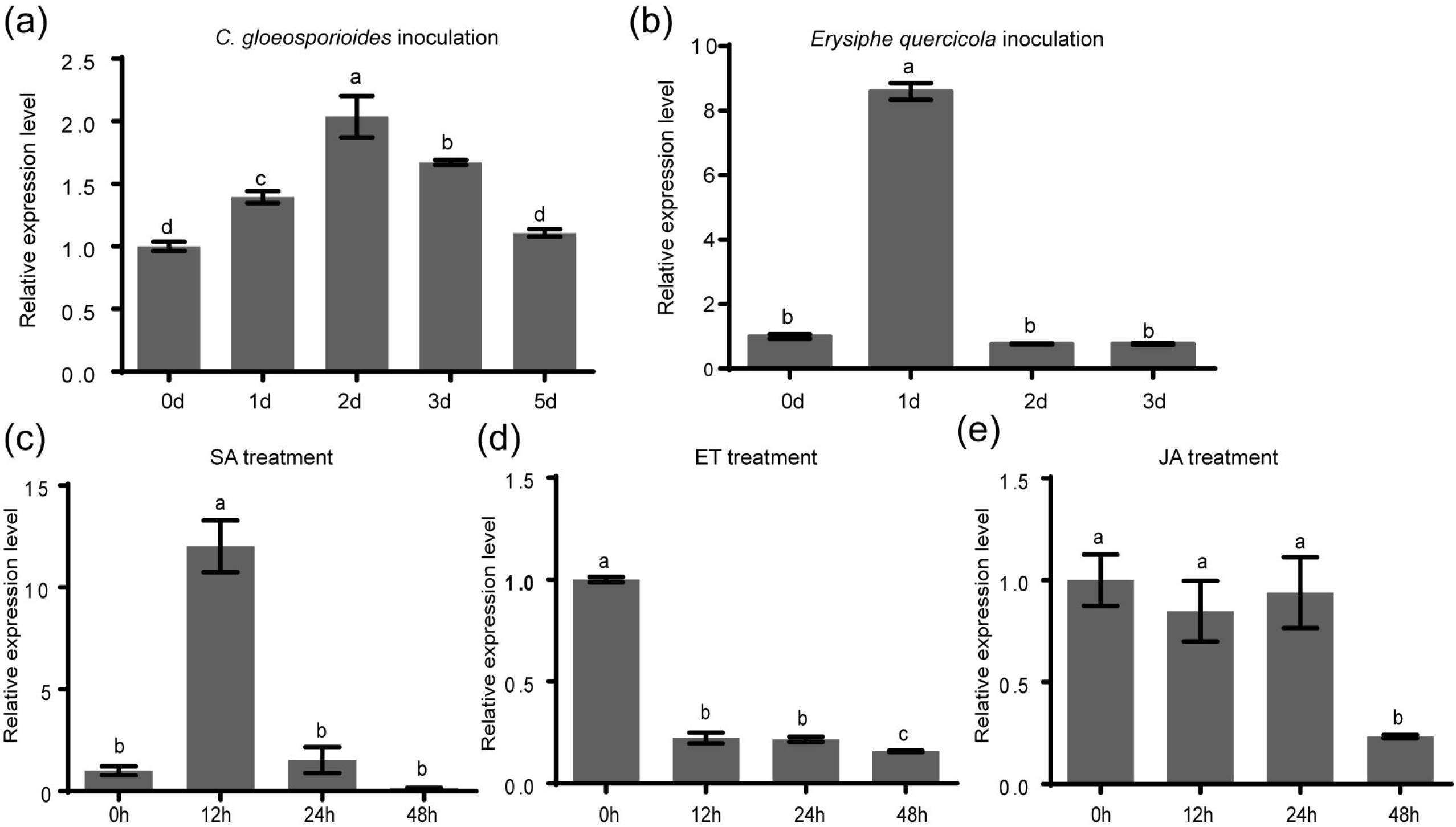
Expression profiles of *HbMYB8-like* in rubber tree the leaves with phathomycete inoculation and phytohormones treatments. Data are shown as the means ± SD from three independent experiments and columns with different letters indicate significant difference (P < 0.05).

### CgNLP1 disrupted the nuclear accumulation of HbMYB8 and inhibited HbMYB8-like induced cell death

We had demonstrated that HbMYB8-like localized on the nucleus as a transcriptional factor and induced necrosis in tobacco tissues (Figure 6). When we transiently co-expressed CgNLP1-GFP fusion and nucleo-scytoplasmic marker (MIEL1-RFP) in *N. benthamiana* leaves, the CgNLP1-GFP completely overlaps with the MIEL1-RFP, indicating that CgNLP1 was localized on the nucleus and cell membrane. To determine whether CgNLP1 interfered the subcellular localization of HbMYB8-like, HbMYB8-like-RFP and CgNLP1-GFP were transiently co-expressed in *N. benthamiana* leaves. Co-expression of HbMYB8-like-RFP with CgNLP1-GFP caused a distinct change in the localization of HbMYB8-like-RFP in *N. benthamiana* cells. By itself, HbMYB8-like-RFP was rather uniformly distributed in the nuclei. However, in the presence of CgNLP1, less HbMYB8-like-RFP was observed in nuclei, and instead a substantial amount of HbMYB8-like was localized on cell membrane. CgNLP1-GFP was also detected on the nucleus and cell membrane (Figure 7a). In addition, we observed that the size of necrosis caused by co-expression of HbMYB8-like-RFP with CgNLP1-GFP was significant smaller than that caused by expression of HbMYB8-like-RFP (Figure 7b and 7c). These data suggested that CgNLP1 could interfere in the nuclear accumulation of HbMYB8-like to inhibit HbMYB8-like induced necrosis.

**Fig.7.**
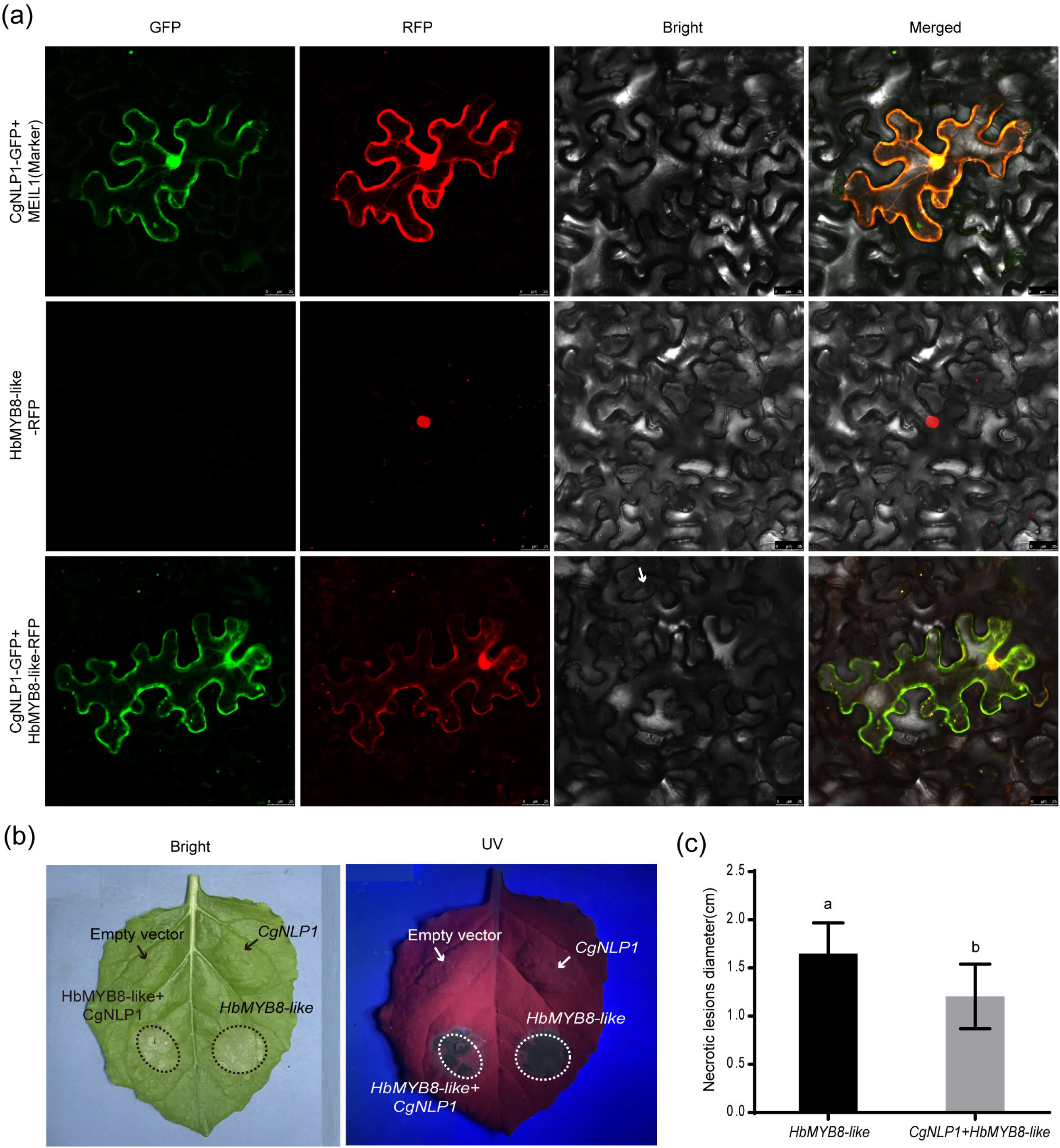
CgNLP1 repressed nuclear accumulation of HbMYB8-like and inhibited necrosis induced by HbMYB8-like. (a) CgNLP1 repressed nuclear accumulation of HbMYB8-like. Scale bar□=□25um. (b) CgNLP1 inhibited necrosis induced by HbMYB8-like. (c) Statistical analysis of necrosis diameter in tobacco leaves. Data are shown as the means ± SD from three independent experiments and columns with different letters indicate significant difference (P < 0.05).

### CgNLP1 inhibited SA signaling mediated by HbMYB8-like in tobacco leaves

Since CgNLP1 could induce ethylene production in plants and HbMYB8 was up-regulated in response to exogenous SA (Figure 2c and Figure 6), we examined the expression patterns of genes related to SA signaling pathway in the tobacco leaves expressing HbMYB8-like and co-expressing HbMYB8-like and CgNLP1. As showed in Figure 8, the expression of *NtPR1a, NtPR1b, NtNPR1* and *NtPAL1* had been enhanced significantly in the tobacco leaves expressing HbMYB8-like for 24 hours, but reduced significantly in the tobacco leaves co-expressing HbMYB8-like and CgNLP1. These results indicated that HbMYB8-like promoted SA signaling and CgNLP1 repressed SA signaling mediated by HbMYB8-like.

**Fig. 8.**
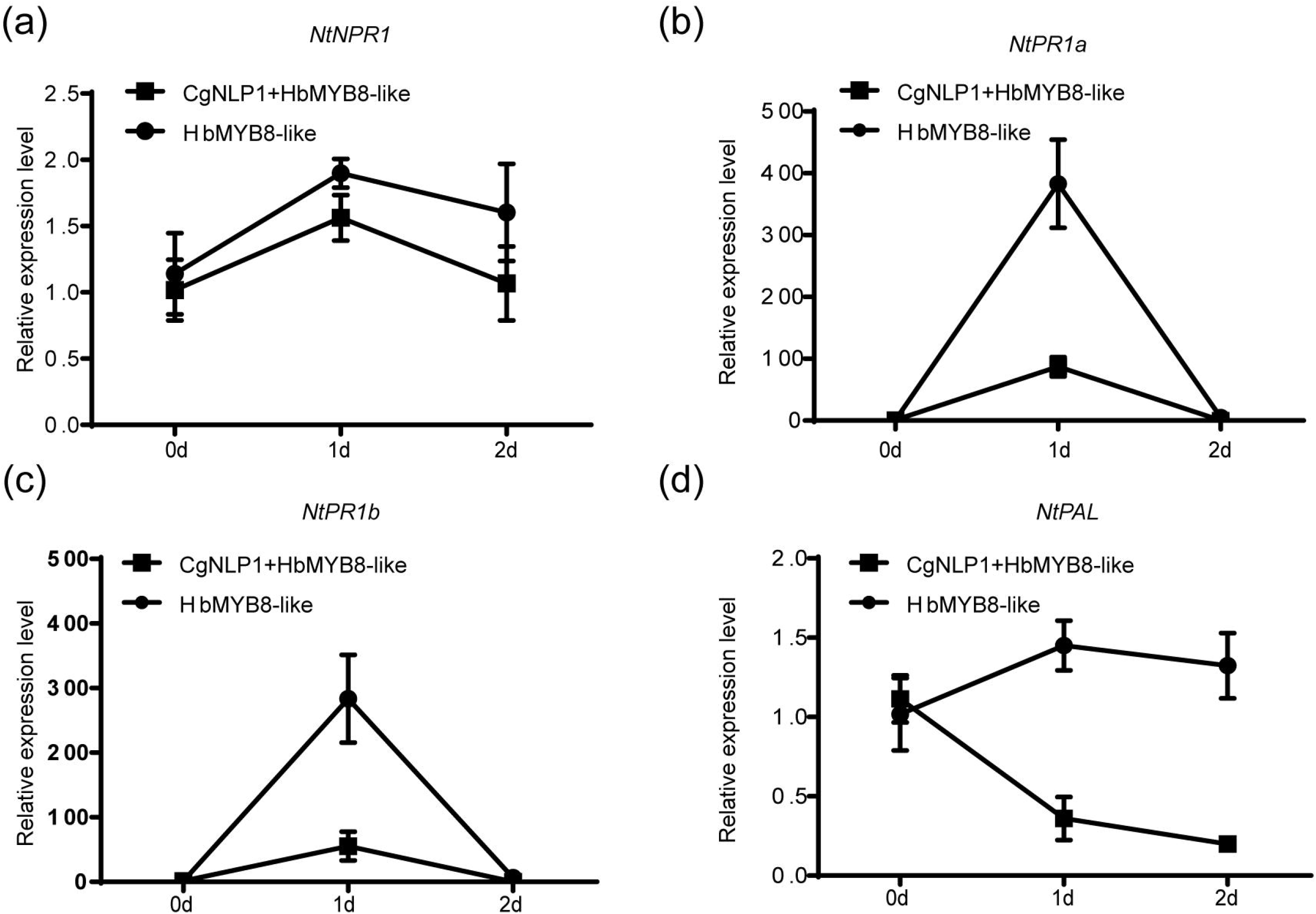
Expression patterns of genes related to SA signaling pathway in the tobacco leaves expressing HbMYB8-like and co-expressing HbMYB8-like and CgNLP1. Data are shown as the means ± SD from three independent experiments.

## Discussion

In the 30 years since the first NLP protein was discovered from the culture filtrate of *Fusarium oxysporum* f.sp. *erythroxyli* (Bailey, 1995), a large number of NLPs had been identified based on a prominent feature of the NPP1 domain with a conserved heptapeptide motif GHRHDWE (Oome & Van den Ackerveken, 2014). As mentioned in the introduction, NLP could be divided into three types, which differed especially in the number of cysteines. TypeINLPs contained two cysteines forming a conserved disulfide bond and typeIIhad four cysteines forming two conserved disulfide bond and an additional putative calcium-binding domain (Oome et al., 2014). In our study, CgNLP1, an NLP protein identified from *C. gloeosporioides* vs. *hevea brasiliensis*, contained a NPP1 domain with two cysteine residues (Figure 1a), which conformed to the structural characteristics of typeINLPs. So, we identified it as type INLP protein and it was also supported by phylogenetic analysis (Figure 1b). It had been suggested that signal peptide at N-terminal, two or four cysteine residues and heptapeptide motif GHRHDWE were directly related to the necrosis inducing ability of NLPs ( Zaparoli et al., 2011; Lenarčič et al., 2017). When one cysteine was replaced, the NLP protein from *Phytophthora parasitica*, NPP1, lost necrosis inducing ability (Fellbrich et al., 2002). Both NLPPp and NLPPs, the NLPs from *P. parasitica* and *P. sojae* respectively, without signal peptide did not show cytotoxicity when transiently expressed in sugar beet (Qutob et al., 2006). In conserved heptapeptide motif GHRHDWE of NLP from *Moniliophthora perniciosa*, the substitution of either at His2 (H) or Asp4 (D) with alanine substantially abrogated its necrosis inducing activity (Ottmann et al., 2009; Zaparoli et al., 2011). However, HaNLP3, an NLP protein from *H. arabidopsidis*, did not induce necrosis on plant cells despite having a conserved heptapeptide motif (Ottmann et al., 2009; Cabral et al., 2012). In our case, CgNLP1 contained a signal peptide, two cysteine residues and an SHRHDWE motif with His2 (H) and Asp4 (D), despite which was different from the typic heptapeptide motif GHRHDWE at the first amino acid site. However, CgNLP1 did not induce obvious necrosis and cell death when transiently expressed in tobacco leaves (Figure S3), suggested that signal peptide, the number of cysteine residue and the conserved SHRHDWE motif were not required for NLPs to induce necrosis and cell death. The mechanism of NLP induced necrosis requires more researches.

NLPs were suggested to play dual roles in plant pathogen interactions as virulence factors and as triggers of plant innate immune responses (Qutob et al., 2006). The deletion strains of NLP1 and NLP2 from *Verticillium dahlia* were found to be significantly less pathogenic on tomato plants (Santhanam et al., 2013). Gene silencing analysis showed that three NLPs from *P. capsica*, PcNLP2, PcNLP6, and PcNLP14, contributed positively to virulence on host plants (Feng et al., 2014). Introduction of Nep1 from *F. oxysporum* into a hypovirulent *Colletotrichum coccodes* strain dramatically increased its virulence and expanded its host spectrum (Amsellem et al., 2002). These data showed that some NLPs were required for virulence of phytopathogens. On the other hand, as described in the introduction part, some NLPs induced plant innate immune responses resulting in enhanced disease resistance (Seidl & Van den Ackerveken, 2019). For example, Ectopic expression of HaNLP3 in Arabidopsis enhanced the resistance to downy mildew (Oome et al., 2014). Our results showed that the disruption of *CgNLP1* impaired the pathogenicity of *C. gloeosporioides* to rubber tree, and ectopic expression of CgNLP1 in Arabidopsis enhanced the resistance to *Botrytis cinerea* and *Alternaria brassicicola* (Figures 3), indicating that *CgNLP1* performed dual functions as both virulence factor and elicitor. Hormone signaling crosstalk played major roles in plant defense against pathogens (Derksena, et al., 2013). SA, JA and ET play critical roles in the regulation of signaling networks of basal resistance against multiple pathogens (Pieterse et al., 2012). SA signaling positively induces plant defense against biotrophic pathogens, whereas the JA/ET pathways are required for resistance predominantly against necrotrophic pathogens and herbivores insects (Yang et al., 2015). We had showed that CgNLP1could enhance disease resistance of Arabidopsis to necrotrophic fungi *B. cinerea* and *A. brassicicola* (Figure 3), and had also detected that CgNLP1 promoted ethylene synthesis and accumulation in plant tissues (Figure 2c), indicating that the disease resistance triggered by CgNLP1 to necrotrophic pathogens may be related to ethylene accumulation and enhanced ethylene mediated signaling pathway.

Fungal effectors were usually secreted and delivered into host plants as pathogenic factors to shield the fungus, suppress the host immune response, or manipulate host cell physiology through targeting plant defense components, signaling, and metabolic pathways to promote host plant colonization (Lo Presti et al., 2015). Plant transcription factors (TFs) played roles in diverse biological processes including defense responses to pathogens (Seo & Choi, 2015). Thus, plant transcription factors are logical targets for effectors, and several studies had demonstrated this. A bacterial effector AvrRps4 interacted with WRKY transcription factors (Sarris et al., 2015). A *Phytophthora infestans* effector Pi03192 prevented the translocation of NAC transcription factors from the endoplasmic reticulum to the nucleus (Hazel et al., 2013). PpEC23 from *Phakopsora pachyrhizi* targeted transcription factor GmSPL12l to suppress host defense response (Qi et al., 2016). Stripe rust effector PstGSRE1 disrupts nuclear localization of transcription factor TaLOL2 to defeat ROS-induced defense in wheat (Qi et al., 2019). In our study, CgNLP1 physically interacted with HbMYB8-like, a R2R3 type MYB transcription factor in rubber tree, which was localized to the nucleus (Figure 5a). The expression of CgNLP1 in tobacco leaves induced partial re-localization of HbMYB8-like from nucleus to plasma membrane and reduced the amount of HbMYB8-like in nuclei (Figure 7a). Logically, it was not necessary for CgNPL1 to make the long journey from the nucleus to bring HbMYB8-like back to the plasma membrane if only to inhibit the function of HbMYB8-like, so it remained to determine the physiological function of HbMYB8-like re-localization.

SA was believed to play important roles in the regulation of programed cell death or hypersensitive reaction in plants, a form of plant defense (Radojičić et al., 2018). According to our data, CgNLP1 contributed to the virulence of *C. gloeosporioides* on rubber tree (Figure 2a and 2b), and its target in rubber tree, HbMYB8-like, induced typic necrosis and cell death (Figure 5b), moreover, CgNPL1 repressed cell death induced by HbMYB8-like in tobacco leaves (Figure 7b and 7c). In addition, it also showed that HbMYB8-like enhanced the expression of genes related to SA signaling pathway such as *NtPR1a, NtPR1b, NtNPR1* and *NtPAL1,* which were inhibited significantly when CgNPL1 co-expressed with HbMYB8-like (Figure 8). Consider the above data, it looked like that the pathogenicity of CgNLP1 was achieved through inhibition of defense induced by HbMYB8-like mediated SA signaling.

In summary, we identified an important *C. gloeosporioides* effector CgNLP1 which played dual roles as virulence factor to rubber tree and as a trigger of plant defense responses, and CgNLP1 blocked nuclear accumulation of rubber tree transcription factor HbMYB8-like and repressed SA signaling mediated by HbMYB8-like. This extended our knowledge of novel pathogenic strategy mediated by effector CgNLP1 for *C. gloeosporioides* on rubber tree.

## Acknowledgements

This study was supported by grants from Hainan Natural Science Foundation (Project No. 319QN167) and the National Natural Science Foundation of China (Grant Nos. 31860478 and 32060591).

## Supporting Information

Additional supporting information may be found in the online version of this article.

**Fig. S1** Nucleotide sequence and deduced amino acid sequence of *CgNLP1*. Shading indicates the amino acid sequences of signal peptide.

**Fig. S2** Generation and molecular confirmation of *CgNLP1* deletion mutant (Δ*CgNLP1*) and complementation mutant (Res-ΔCg*NLP1*). (a) The diagram of *CgNLP1* knockout vector. (b) Diagnostic PCR analysis for deletion of *CgNLP1* and integration of *CgNLP1* into the genome of *C. gloeosporioides*. (c) The expression level of *CgNLP1* in WT, Δ*CgNLP1* and Res-Δ*CgNLP1* by quantitative RT-PCR.

**Fig. S3** Trypan blue transient of tobacco leaves expressing *CgNLP1*. pEGAD indicates the tissue expressing empty vector pEGAD-eGFP, pEGAD-CgNLP1 indicates the tissue expressing recombinant vector pEGAD-*CgNLP1*-eGFP.

**Fig. S4** Identification of *CgNLP1* transgenic Arabidopsis lines by Western blot. Col-0 indicated Arabidopsis Columbia-0 and 1-6 indicated different CgNLP1-OE transgenic lines under Col-0 background.

**Fig. S5** Alignment of HbMYB8-like and homologs from different plants. Shading indicated regions of conservation in all (black), the same amino acid as HbMYB8-like (grey) of sequences. The protein sequences used for alignment are: *Arabidopsis thaliana* AtMYB6 (EFH48703.1), *Arabidopsis thaliana* AtMYB8 (Q9SDS8.1), *Oryza sativa* OsMYB30 (Q6K1S6.1), *Gossypium hirsutum* GhMYB1 (NP_001313761.1), *Triticum aestivum* TaRIM1 (AMP18876.1).

**Fig S6** Phylogenetic tree of HbMYB8-like with different types of MYB proteins in plants.

**Table S1** The primers used for PCR amplification quantitative and RT-PCR.

## Refferences

Albert I, Böhm H, Albert M, Feiler EC, Imkampe J, Wallmeroth N, Brancato C, Raaymakers MT, Oome S, Zhang H et al. 2015. An RLP23-SOBIR1-BAK1 complex mediates NLP-triggered immunity. Nature Plants 1:15140.

Alkan N, Friedlander G, Ment D, Prusky D, and Fluhr, R. 2015. Simultaneous transcriptome analysis of *Colletotrichum gloeosporioides* and tomato fruit pathosystem reveals novel fungal pathogenicity and fruit defense strategies. New Phytologist 205: 801–815.

Amsellem Z, Cohen BA, Gressel J. 2002. Engineering hypervirulence in a mycoherbicidal fungus for efficient weed control. Nature Biotechnology 20:1035–1039.

Azmi NA, Singkaravanit-Ogawa S, Ikeda K, Kitakura S, Inoue Y, Narusaka Y, Shirasu K, Kaido M, Mise K, Takano Y. 2018. Inappropriate expression of an nlp effector in *Colletotrichum orbiculare* impairs infection on *Cucurbitaceae Cultivars* via plant recognition of the c-terminal region. Molecular Plant-microbe Interactions 31: 101–111.

Bailey BA. 1995. Purification of a protein from culture filtrates of *Fusarium oxysporum* that induces ethylene and necrosis in leaves of *Erythroxylum coca*. Phytopathology 85:1250–1255.

Böhm H, Albert I, Oome S, Raaymakers TM, Van den Ackerveken G, Nürnberger T. 2014. A conserved peptide pattern from a widespread microbial virulence factor triggers pattern-induced immunity in Arabidopsis. PLoS Pathogens 10: e1004491.

Cabral A, Oome S, Sander N, Küfner I, Nürnberger T, Van den Ackerveken G. 2012. Nontoxic Nep1-like proteins of the downy mildew pathogen *Hyaloperonospora arabidopsidis:* repression of necrosis-inducing activity by a surface-exposed region. Molecular Plant-microbe Interactions 25: 697–708.

Chen J, Bao S, Fang Y, Wei L, Zhu W, Peng Y, Fan J. 2021. An LRR-only protein promotes NLP-triggered cell death and disease susceptibility by facilitating oligomerization of NLP in Arabidopsis. New Phytologist. doi:10.1111/nph.17680.

Chen X, Huang S, Zhang Y, Gui L, Li Y, Zhu F. 2018. Identification and functional analysis of the NLP-encoding genes from the phytopathogenic oomycete *Phytophthora capsici*. Molecular Genetics and Genomics 293:931–943.

Derksena H, Rampitschb C, Daayf F. 2013. Signaling cross-talk in plant disease resistance. Plant Science 207: 79–87.

Dodds PN, Rathjen JP. 2010. Plant immunity: towards an integrated view of plant-pathogen interactions. Nature Reviews Genetics 11: 539–548.

Dong S, Kong G, Qutob D, Yu X, Tang J, Kang J, Dai T, Wang H, Gijzen M, Wang Y. 2012. The NLP toxin family in *Phytophthora sojae* includes rapidly evolving groups that lack necrosis-inducing activity. Molecular Plant-microbe Interactions 25: 896–909.

Seo E, Choi D. 2015. Functional studies of transcription factors involved in plant defenses in the genomics era. Briefings in functional genomics 14: 260–267.

Fang Y, Peng Y, Fan J. 2017. The Nep1-like protein family of *Magnaporthe oryzae* is dispensable for the infection of rice plants. Scientific Reports 7: 4372.

Fellbrich G, Blume B, Brunner F, Hirt H, Kroj T, and Ligterink W, Romanski A, Nürnberger T. 2000. *Phytophthora parasitica* elicitor-induced reactions in cells of *Petroselinum crispum*. Plant and Cell Physiology 41: 692–701.

Fellbrich G, Romanski A, Varet A, Blume B, Brunner F, Engelhardt S, Felix G, Kemmerling B, Krzymowska M, Nürnberger T. 2002. NPP1, a Phytophthora-associated trigger of plant defense in parsley and Arabidopsis. The Plant Journal 32: 375–390.

Feng B, Zhu X, Fu L, Lv R, Storey D, Tooley P, Zhang X. 2014. Characterization of necrosis-inducing NLP proteins in *Phytophthora capsici*. BMC Plant Biology 14:126

Gijzen M, Nürnberger T. 2006. Nep1-like proteins from plant pathogens: recruitment and diversification of the npp1 domain across taxa. Phytochemistry, 67(16), 1800–1807.

Guy E, Lautier M, Chabannes M, Roux B, Lauber E, Arlat M, Noël LD. 2013. XopAC-triggered immunity against xanthomonas depends on arabidopsis receptor-like cytoplasmic kinase genes *PBL2* and *RIPK*. Plos One 8: e73469.

Hazel M, Petra C, Boevink MR, Armstrong LP, Sonia G, Juan M, Stephen CW, Jim LB, Paul RJ, Birch M. 2013. An RxLR effector from *Phytophthora infestans* prevents re-localisation of two plant NAC transcription factors from the endoplasmic reticulum to the nucleus. PLoS Pathogens 9: e1003670.

Heard S, Brown NA, Hammond-Kosack K. 2015. An Interspecies comparative analysis of the predicted secretomes of the necrotrophic plant pathogens *Sclerotinia sclerotiorum* and *Botrytis cinerea*. PLoS One 10: e0130534.

Kanneganti TD, Huitema E, Cakir C, Kamoun S. 2006. Synergistic interactions of the plant cell death pathways induced by *Phytophthora infestans* Nepl-like protein PiNPP1.1 and INF1 elicitin. Molecular Plant-microbe Interactions 19:854–63

Kleemann J, Rincon-Rivera LJ, Takahara H, Neumann U, van Themaat EVL, van der Does HC, Hacquard S, Stüber K, Will I, Schmalenbach W et al. 2012. Sequential delivery of host-induced virulence effectors by appressoria and intracellular hyphae of the phytopathogen *Colletotrichum higginsianum*. PLoS Pathogens. 8: e1002643.

Lenarčič T, Albert I, Böhm H, Hodnik V, Pirc K, Zavec AB, Podobnik M, Pahovnik D, Žagar E, Pruitt R et al. 2017. Eudicot plant-specific sphingolipids determine host selectivity of microbial NLP cytolysins. Science 358: 1431.

Levin E, Raphael G, Ma J, Ballester AR, Feygenberg O, Norelli J, Aly R, Gonzalez-Candelas L, Wisniewski M, Droby S. 2019. Identification and functional analysis of NLP-encoding genes from the postharvest pathogen *Penicillium expansum*. Microorganisms 7:175.

Liang C, Zhang B, Zhou Y, Yin H, An B, Lin D, He C and Luo H. 2021. CgNPG1 as a novel pathogenic gene of *Colletotrichum gloeosporioides* from *Hevea brasiliensis* in mycelial growth, conidiation, and the invasive structures development. Frontiers in Microbiology 12:629387.

Liu X, Li B, Cai J, Zheng X, Feng Y, and Huang G. 2018. *Colletotrichum* species causing anthracnose of rubber trees in china. Scientific Reports 8:10435.

Lo Presti L, Lanver D, Schweizer G, Tanaka S, Liang L, Tollot M, Zuccaro A, Reissmann S, Kahmann1 R. 2015. Fungal effectors and plant susceptibility. Annual Review of Plant Biology 66:513–545.

Oome S, Van den Ackerveken G. 2014. Comparative and functional analysis of the widely occurring family of Nep1-like proteins. Molecular Plant-Microbe Interactions 27:1081–1094.

Oome S, Raaymakers TM, Cabral A, Samwel S, Bohm H, Albertc I, Nürnbergerc T, Van den Ackervekena G. 2014. Nep1-like proteins from three kingdoms of life act as a microbe-associated molecular pattern in Arabidopsis. PNAS 111:16955–60.

Ottmann C, Luberacki B, Küfner I, Koch W, Brunner F, Weyand M, Mattinend L, Pirhonend M, Anderluhe G, Seitza HU et al. 2009. A common toxin fold mediates microbial attack and plant defense. Proceedings of the National Academy of Sciences 106: 10359–10364.

Pieterse CM, Van der Does D, Zamioudis C, Leon-Reyes A, Van Wees SC. 2012. Hormonal modulation of plant immunity. Annual Review of Cell and Developmental Biology 28: 489–521.

Qi MS, Link TI, Müller M, Hirschburger D, Pudake RN, Pedley KF, Braun E, Voegele RT, Baum TJ, Whitham SA. 2012. A small cysteine-rich protein from the Asian soybean rust fungus, *Phakopsora pachyrhizi*, suppresses plant immunity. PLoS Pathogens 12: e1005827.

Qi T, Guo J, Liu P, He F, Wan C, Islam MA, Tyler BM, Kang Z, and Guo J. 2019. Stripe rust effector PstGSRE1 disrupts nuclear localization of ROS-promoting transcription factor TaLOL2 to defeat ROS-induced defense in wheat. Molecular Plant 12: 1624–1638.

Qutob D, Kemmerling B, Brunner F, Küfner I, Engelhardt S, Gust AA, Luberacki B, Seitz HU, Stahl D, Rauhut T et al. 2006. Phytotoxicity and innate immune responses induced by Nep1-like proteins. The Plant Cell 18: 3721–374.

Radojicic A, Li X, Zhang Y. 2018. Salicylic acid: a double-edged sword for programed cell death in plants. Frontiers in Plant Science 9:1133.

Rauhut T, Luberacki B, Seitz HU, Glawischnig E. 2009. Inducible expression of a Nep1-like protein serves as a model trigger system of camalexin biosynthesis. Phytochemistry 70:185–189.

Santhanam P, van Esse HP, Albert I, Faino L, Nürnberger T, Thomma BP. 2013. Evidence for functional diversification within a fungal NEP1-like protein family. Molecular Plant-Microbe Interactions 26: 278–286.

Sarris PF, Duxbury Z, UnHuh S, Ma Y, Segonzac C, Sklenar J, Derbyshire P, Cevik V, Rallapalli G, Saucet, SB et al. 2015. A plant immune receptor detects pathogen effectors that target WRKY transcription factors. Cell 161: 1089–1100.

Schouten A, Baarlen PV, and Kan J. 2008. Phytotoxic Nep1-like proteins from the necrotrophic fungus *Botrytis cinerea* associate with membranes and the nucleus of plant cells. New Phytologist 177: 493–505.

Schumacher S, Grosser K, Voegele RT, Kassemeyer H-H, Fuchs R. 2020. Identification and characterization of Nep1-like proteins from the grapevine downy mildew pathogen *Plasmopara viticola*. Frontiers in Plant Science 11:65.

Seidl MF, Ackerveken G. 2019. Activity and phylogenetics of the broadly occurring family of microbial Nep1-like proteins. Annual Review of Phytopathology 57:1–20.

Van den Ackerveken G. 2017. How plants differ in toxin-sensitivity. Science 358:1383–1384.

Villela-Dias C, Camillo LR, de Oliveira GA, Sena JA, Santiago AS, de Sousaa S, Mendesa JS, Pirovania CP, Alvima F C, Costa MGC. 2014. Nep1-like protein from *Moniliophthora perniciosa* induces a rapid proteome and metabolome reprogramming in cells of *Nicotiana benthamiana*. Physiologia Plantarum 150:1–17

Wang Q, An B, Hou X, Guo Y, Luo H and He C. 2017. Dicer-like proteins regulate the growth, conidiation, and pathogenicity of *Colletotrichum gloeosporioides* from *hevea brasiliensis*. Frontiers in Microbiology 8: 2621.

Yang Y, Ahammed JG, Wu C, Fan S, Zhou Y. 2015. Crosstalk among Jasmonate, Salicylate and Ethylene Signaling Pathways in Plant Disease and Immune Responses. Current Protein and Peptide Science 16: 450–461.

Yang J, Wang Q, Luo H, He, C, An B. 2020. HbWRKY40 plays an important role in the regulation of pathogen resistance in *Hevea brasiliensis*. Plant Cell Reports 39:1095–1107.

Zaparoli G, Barsottini MR, de Oliveira JF, Dyszy F, Teixeira PJ, Barau JG, Garcia O, Costa-Filho AJ, Ambrosio AL, Pereira GA et al. 2011. The crystal structure of necrosis- and ethylene-inducing protein 2 from the causal agent of cacao’s witches’ broom disease reveals key elements for its activity. Biochemistry 50: 9901–9910.

