## Supplementary figures and images for "The effector protein CgNLP1 of *Colletotrichum gloeosporioides* from *Hevea brasiliensis* disrupts nuclear localization of necrosis-induced transcription factor HbMYB8-like to suppress plant defense signaling"

### Figure S1

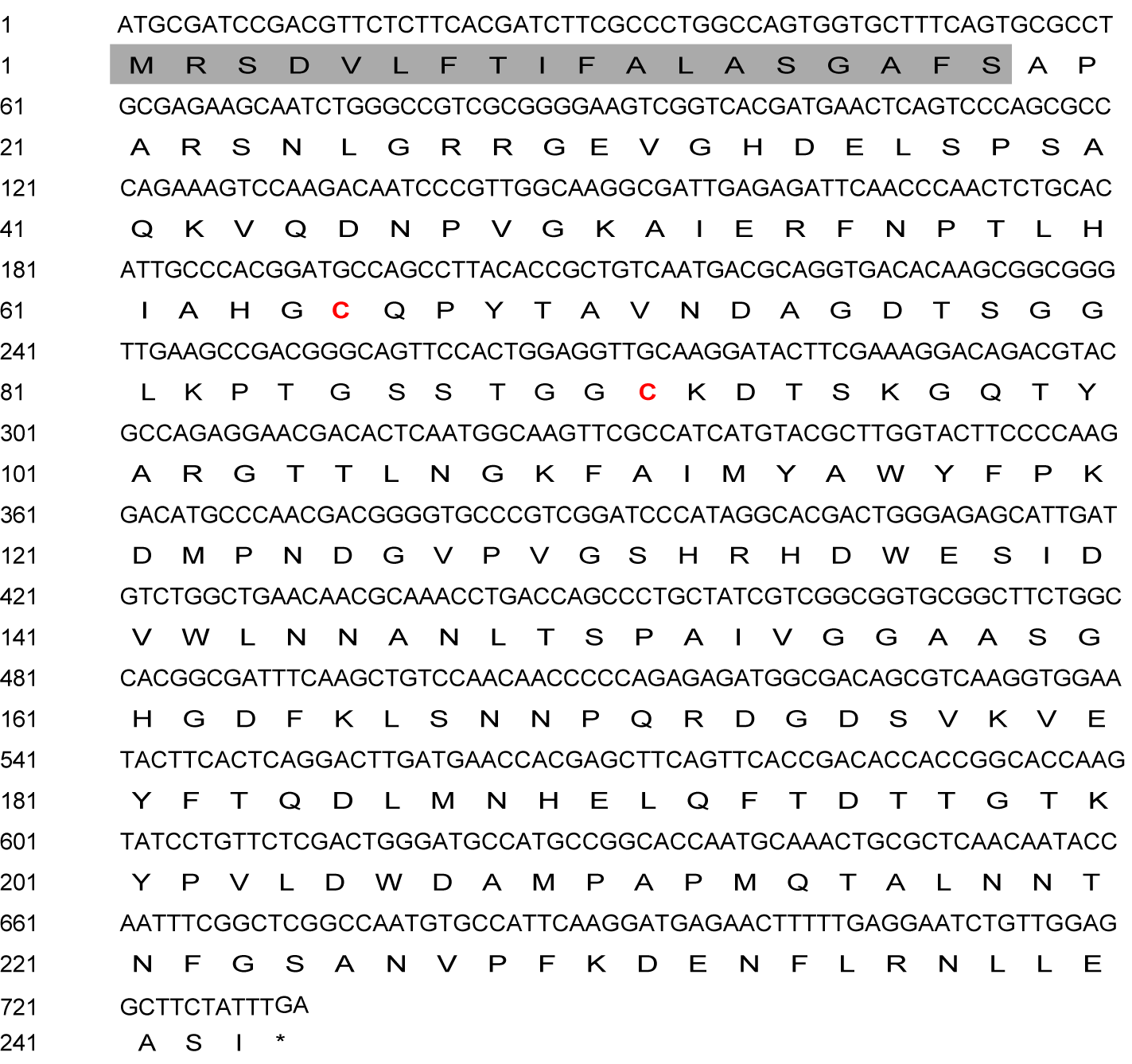

### Figure S2

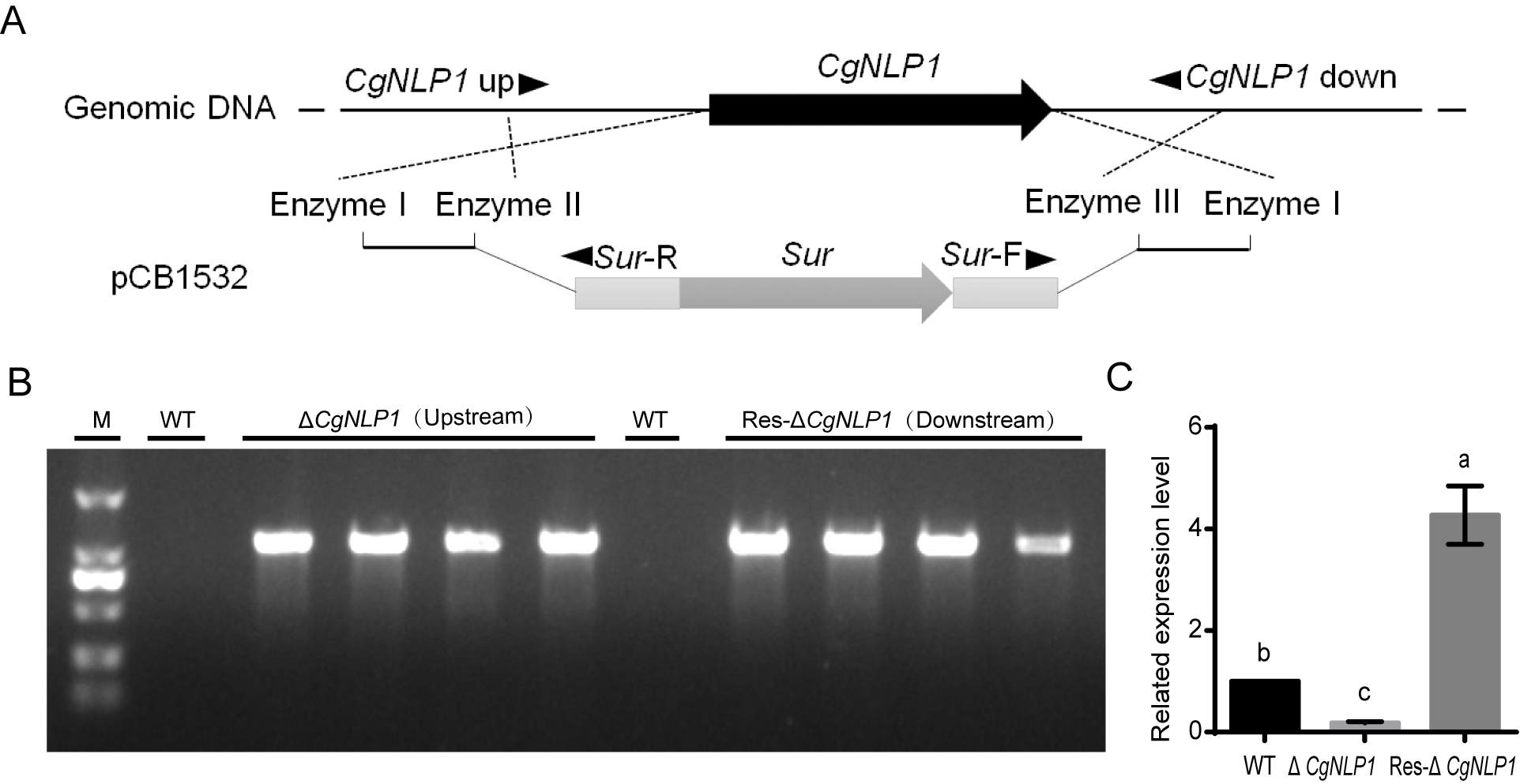

### Figure S3

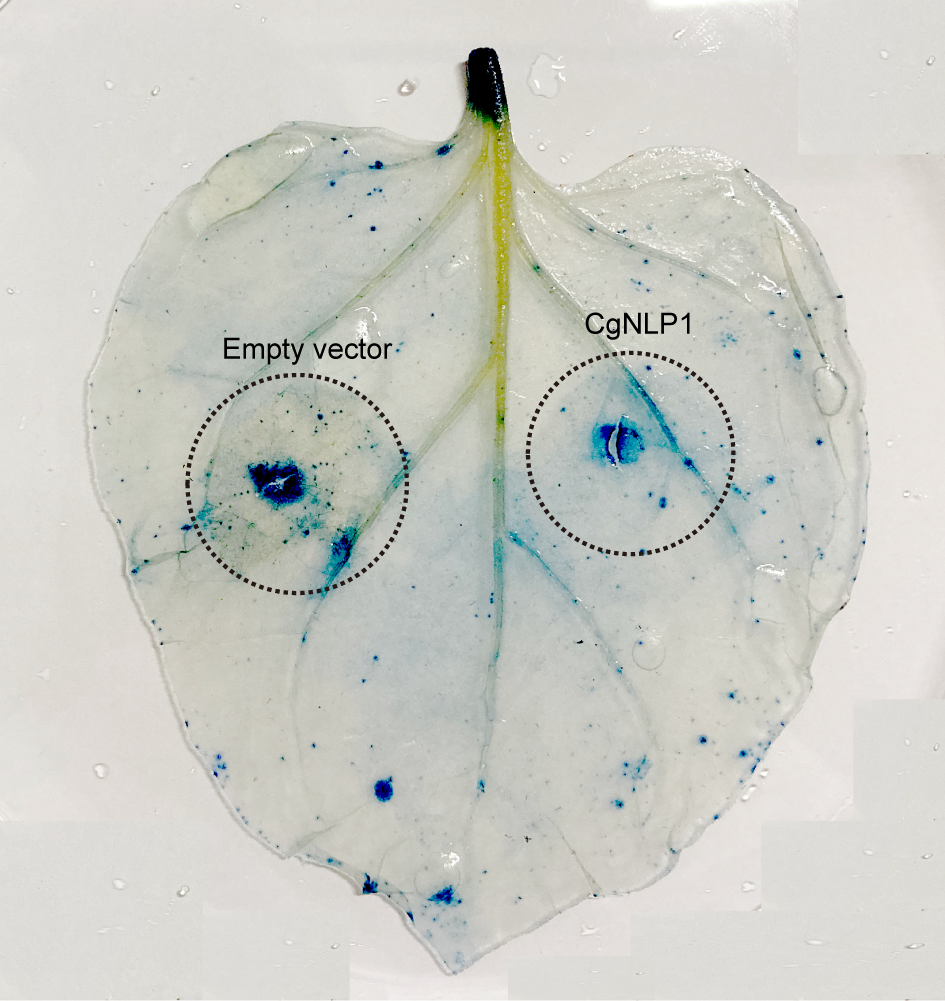

### Figure S4

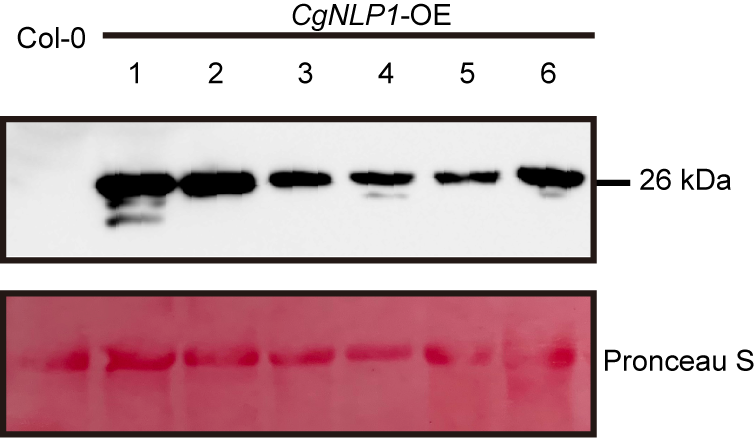

### Figure S5

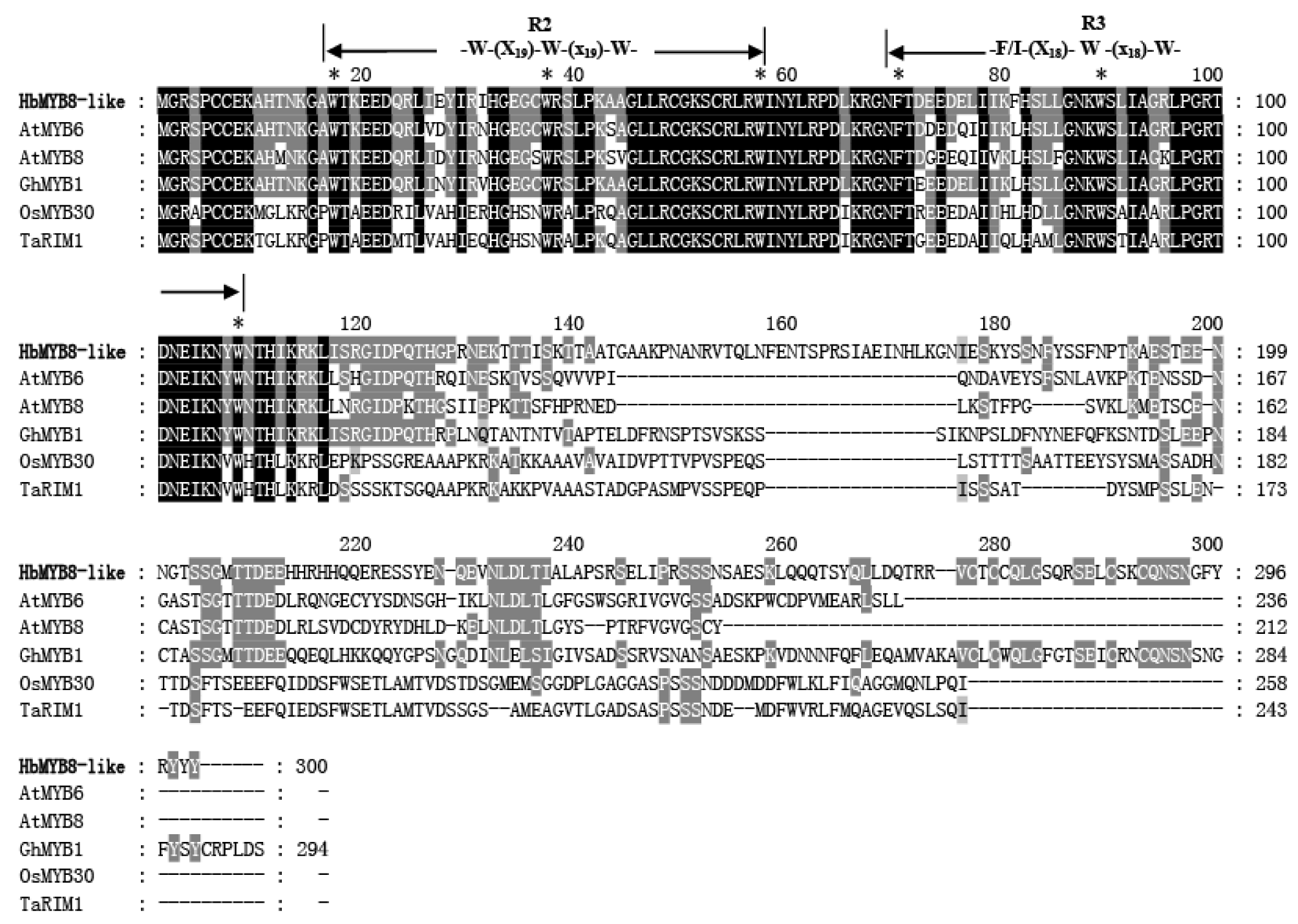

### Figure S6

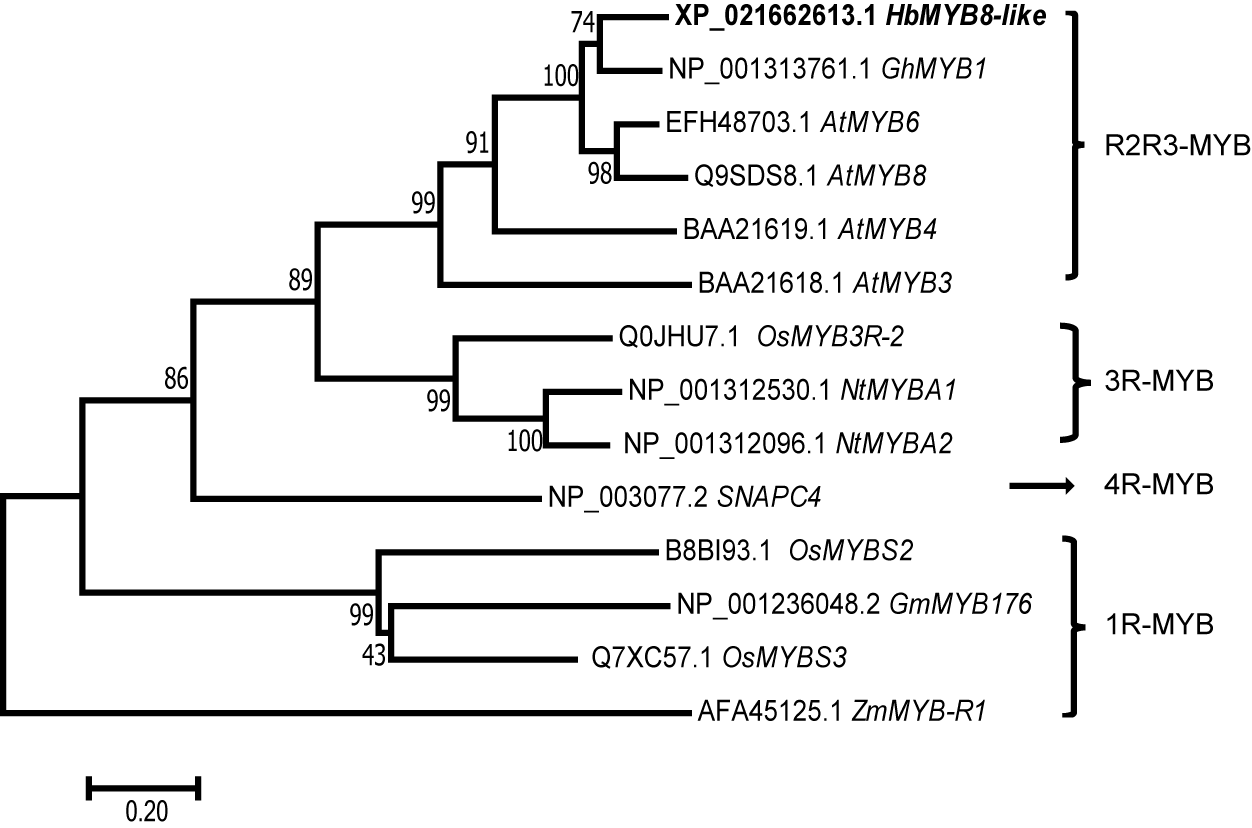
